# The E3 Ubiquitin Ligase Trip12 attenuates Wnt9a/Fzd9b signaling during hematopoietic stem cell development

**DOI:** 10.1101/2024.10.25.620301

**Authors:** Jessica Ensing, Amber D. Ide, Carla Gilliland, Visakuo Tsurho, Isabella Caza, Amber N. Stratman, Nathan J. Lanning, Stephanie Grainger

## Abstract

Wnt signaling is essential for both the development and homeostasis of diverse cellular lineages, including hematopoietic stem cells. Organism-wide, Wnt signals are tightly regulated, as overactivation of the pathway can lead to tumorigenesis. Although numerous Wnt ligands and Frizzled (Fzd) receptors exist, how particular Wnt/Fzd pairings are established and how their signals are regulated is poorly understood. We have previously identified the requirements of the cognate pairing of Wnt9a and Fzd9b for early hematopoietic stem cell proliferation. However, the specific signals governing activation, but equally important, the molecular mechanisms required to turn the signal ‘off,’ are unknown. Here, we show that the E3 ubiquitin ligase Trip12 (thyroid hormone receptor interactor 12) is specifically required to ubiquitinate the third intracellular loop of Fzd9b at K437, targeting it for lysosomal degradation. In contrast to other ubiquitin ligases described to regulate the cell surface availability of multiple Fzds broadly, our data indicate that Trip12 is selective for Fzd9b. We further demonstrate that this occurs through ubiquitination at K437 of Fzd9b in the third intracellular loop, ultimately leading to a decrease in Fzd9b receptor availability and in Wnt9a/Fzd9b signaling that impacts hematopoietic stem cell proliferation in zebrafish. Our results point to specific mechanisms driving the availability of different Fzd receptors. Determining how particular Fzd abundance is regulated at the membrane will be critical to developing specific therapies for human intervention.

**One sentence summary:** Trip12 ubiquitinates Fzd9b

## Introduction

The Wnt signaling pathway is highly conserved among metazoans and drives diverse processes in development and homeostasis, such as tissue patterning, morphogenesis and balancing stem cell proliferation and differentiation^1,2^. The intracellular events in the Wnt pathway are diverse and are typically categorized as either β-catenin-dependent or β-catenin-independent pathways. Here, we focus on β-catenin-dependent signaling. In the b-catenin-dependent pathway, a secreted Wnt protein binds to a seven-pass transmembrane cell surface receptor from the *Frizzled (FZD)* gene family, and a single-pass transmembrane receptor from the *low-density lipoprotein receptor-related protein family* (*LRP5* or *LRP6*). This event at the membrane ultimately leads to dissociation of the cytoplasmic β-catenin destruction complex, enabling translocation of β-catenin to the nucleus for target gene activation and to diverse biological outputs across a multitude of tissue types^1,2^.

The importance of Wnt signaling is reflected by an abundance of driver mutations in Wnt pathway genes, causing diseases such as cancer, Alzheimer’s disease, and diabetes, among others^3–6^. For example, losing mechanisms to downregulate receptor availability at the cell surface can lead to cancer^7–9^. Correspondingly, tight control of signal activation is required at multiple levels, including through the availability of FZD receptors at the membrane. One mechanism to regulate membrane receptor trafficking is post-translational modifications, where amino acid side chains have covalent modifications impacting their function, such as ubiquitination^10^. For example, the E3 Ubiquitin ligases RNF43 and ZNRF3 target FZDs for degradation, a critical step in restricting Wnt signaling^7–10^. Congruently, mutations in genes that negatively regulate membrane localization of FZD, such as RNF43 and ZNRF3, are also common in cancers since loss of function leads to excessive proliferation.

In vertebrates, there are at least 19 Wnts and 10 Fzds, suggesting some degree of functional specificity and ligand-receptor selectivity, although this is only beginning to be uncovered. We recently showed that the specific pairing of Wnt9a and Fzd9(b) is required in zebrafish and human hematopoietic stem cell development^11,12^. Wnt9a/Fzd9b signaling specificity is partially mediated through interaction with the epidermal growth factor receptor (EGFR). Phosphorylation of the Fzd9b tail by EGFR in response to Wnt9a leads to caveolin-mediated endocytosis of the receptor complex, which is required for signal initiation^13^. However, how the membrane availability of Fzd9b is regulated remains incompletely understood. Here, we demonstrate that Fzd9b is ubiquitinated, which is required to attenuate the Wnt9a signal. In addition, we identify that the E3 Ubiquitin ligase TRIP12 is required to restrict Wnt9a/Fzd9b activity by mediating trafficking to the lysosome. Finally, we show that Trip12 plays a role in zebrafish HSC development, a Wnt9a/Fzd9b dependent event.

## Results

### Ubiquitination of Fzd9b at K437 is required for Wnt9a signal attenuation

We have previously reported that Wnt9a and Fzd9b form a cognate pairing required for zebrafish hematopoietic stem cell development^11,12^. Tight regulation of Wnt signals is vital to homeostasis, particularly in stem cell niches; however, regulation of the Wnt9a/Fzd9b signal is incompletely understood. Post-translational modifications such as ubiquitination are essential regulators of receptor trafficking and degradation^10,14^. Here, we sought to identify potential ubiquitin modifications impacting on Fzd9b function.

To do so, we generated a stable human embryonic kidney (HEK)293 cell line harboring an expression construct with zebrafish Fzd9b fused to a V5 tag^15^ and an AviTag^16^ (Fzd9b-V5-AVI) followed by a ribosomal skip sequence (P2A) and the biotinylating enzyme BirA^R118G^ (Fig. 1A, B)^17^ to improve efficacy of precipitating Fzd9b. The Fzd9b-V5-AVI cells stably expressed Fzd9b at levels comparable to our previously established transgenically overexpressed Fzd9b-mKate fusion protein line^13^ (Supplementary Fig. 1A). To ensure that the Fzd9b-V5-AVI was functional, we used an established reporter assay, Super TOP Flash, which responds to Wnt/b-catenin signaling^18^, and found that Fzd9b-V5-AVI was functional in response to Wnt9a (Supplementary Fig. 1B). Further, an exogenous dosage of 0.2 mg/L biotin increased the level of Fzd9b biotinylation *in vivo* (Supplementary Fig. 1C).

**Figure 1:**
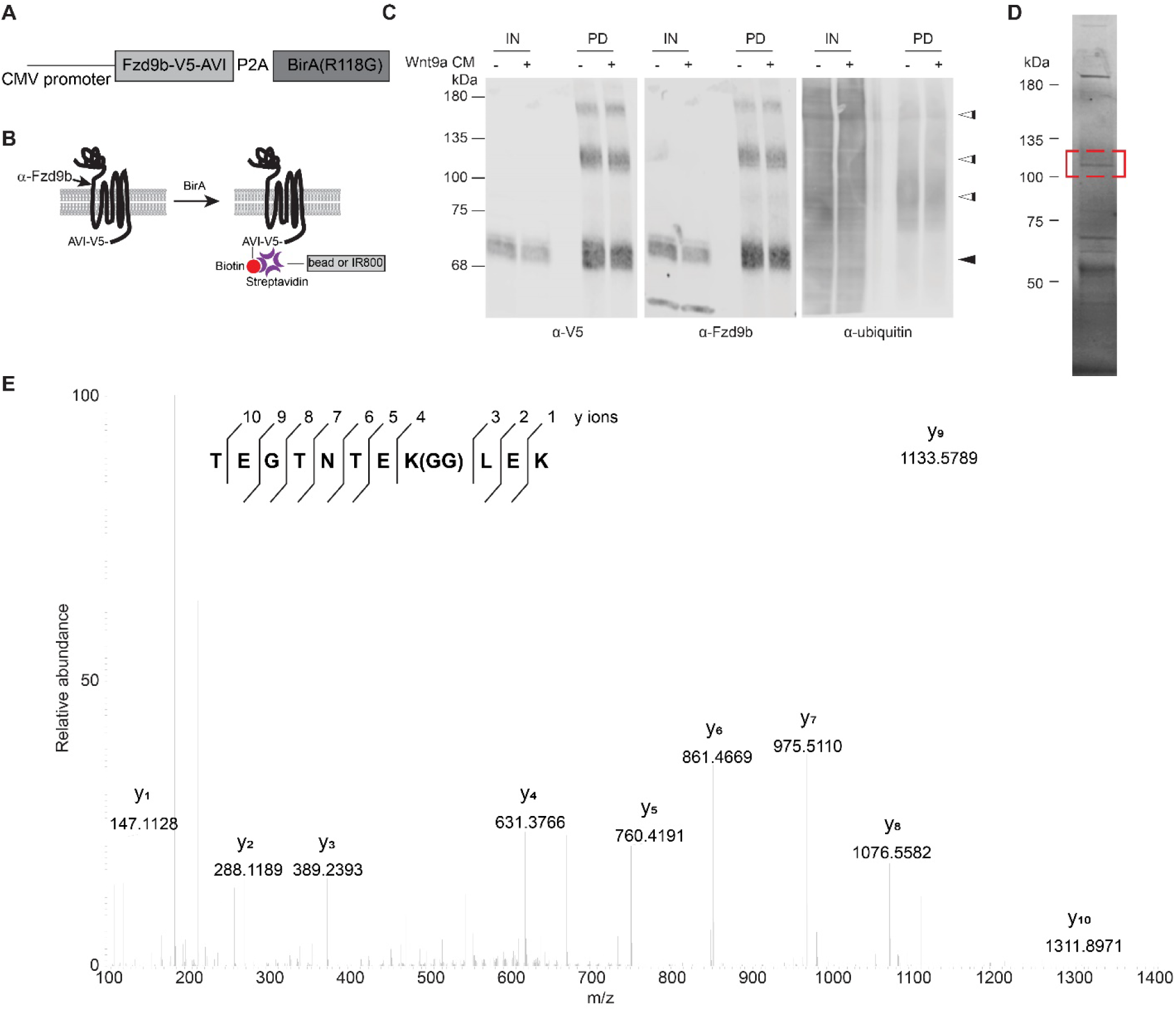
Fzd9b is ubiquitinated. **A.** Schematic of Fzd9b-V5-AVI expression vector. **B.** Fzd9b-V5-AVI is co-expressed with BirA, which biotinylates the AVI tag on Fzd9b, enabling subsequent purification using streptavidin beads, and/or detection using Streptavidin protein. The epitope region for the Fzd9b antibody is noted. **C.** Immunoblot of input (5%, IN) and pulldown (PD) of protein extracted from HEK293 Fzd9b-V5-AVI cells treated with mock or Wnt9a conditioned medium (CM) for one minute. Blots for V5, Fzd9b and ubiquitin antibodies are shown. Black arrowhead points to expected size for Fzd9b; white arrowheads point to additional sized bands detected by V5 and Fzd9b antibodies in PD lanes. **D.** Fzd9b-V5-AVI protein purified by streptavidin beads and stained using Coomassie Blue dye. Band extracted for mass spectrometry analysis is denoted by the red box. **E.** MS2 spectra of the endogenous ubiquitinated peptide (TEGTNTEK(GG)LEK) of Fzb9b, displaying the relative abundance of the fragment ions. All fragment ions were manually assigned. GG denotes the di-glycyl remnant produced on ubiquitinated lysine residues (K-ε-GG) following trypsin digestion.

In the absence of pulldown, we noted a faint, higher molecular weight band around 110 kDa (white arrowhead, Supplementary Fig. 1A) in addition to the expected size of Fzd9b (black arrowhead, Supplementary Fig. 1A), suggesting potential post-translational modifications with larger molecular weights such as the addition of multiple 8kDa ubiquitin moieties. To determine if Fzd9b was modified, and if Wnt9a stimulation changed the status of ubiquitination, we treated

Fzd9b-V5-AVI cells with conditioned medium collected from Chinese hamster ovary (CHO) cells expressing Wnt9a, or naïve CHO cells (mock) for 1-minute, precipitated Fzd9b-V5-AVI using streptavidin beads, and assessed pulldown (PD) fractions using antibodies to V5 and Fzd9b. Using this approach, we observed multiple Fzd9b/V5 bands at the predicted sizes around 70 kDa (Fig. 1C, black arrowhead)^11^. We also noted the enrichment of several bands at larger molecular weights at approximately 85, 110, and 150 kDa (Fig. 1C, white arrowheads), further indicative of putative Fzd9b post-translational modifications and/or formation of protein complexes. We did not note any major changes in modified or unmodified Fzd9b abundance with Wnt9a treatment (Fig. 1C). We further noted that pulldown is required to see many of the modified Fzd9b bands, suggesting that these represent a small portion of the Fzd9b pool (Fig. 1C). To determine if these multiple Fzd9b moieties may represent ubiquitinated receptor, we immunoblotted for total ubiquitin, and found that ubiquitin was enriched at the 85, 110 and 150 kDa band (Fig. 1C, white arrowheads), suggesting that Fzd9b could be ubiquitinated. To test if longer treatments of Wnt9a led to changes in enrichment of the ubiquitinated form of Fzd9b, we treated cells for 25 or 50 minutes, and found that there was a slight increase in ubiquitinated Fzd9b, particularly at 110 kDa (Supplementary Fig. 1D). To confirm that there were ubiquitin moieties specifically attached to Fzd9b, we precipitated Fzd9b from untreated cells, extracted the band around 110 kDa (Fig. 1D), and used mass spectrometry to identify peptides with a signature C-terminal glycine-glycine moiety, indicative of ubiquitination^19^. Using this approach, we identified the peptide TEGTNTEK(GG)LEK, corresponding to a site of ubiquitination at K437 of Fzd9b (Fig. 1E). We validated this using a synthetic peptide, and found identical results (Supplementary Fig. 1E), indicating that Fzd9b is ubiquitinated at K437.

The ubiquitination site at K437 is in the third intracellular loop of Fzd9b (Fig. 2A). There is a very high degree of conservation in the amino acid sequence surrounding K437 among vertebrates (Fig. 2B), implicating the importance of this residue. We tested the impact of losing the ubiquitination site at K437 on Wnt9a/Fzd9b signaling using a Super TOP Flash assay comparing K437R (Arginine (R) is structurally similar to Lysine (K), but cannot be ubiquitinated) to WT Fzd9b, and showed close to a 3-fold increase in Wnt activity in Fzd9b^K437R^ (Fig. 2C), indicating that this residue plays a role in limiting Wnt9a signaling activity, independent of protein levels of Fzd9b (Supplementary Fig. 2). To test if this ubiquitination site plays a conserved role in human WNT9A and FZD9 signaling, we tested the signaling capacity of FZD9^K440R^ in a Super TOP Flash assay and found an analogous increase in signaling (Fig. 2D), indicating that function of this residue is conserved and supporting the notion that K437 (or K440 in humans) delimits the Wnt9a signal.

**Figure 2:**
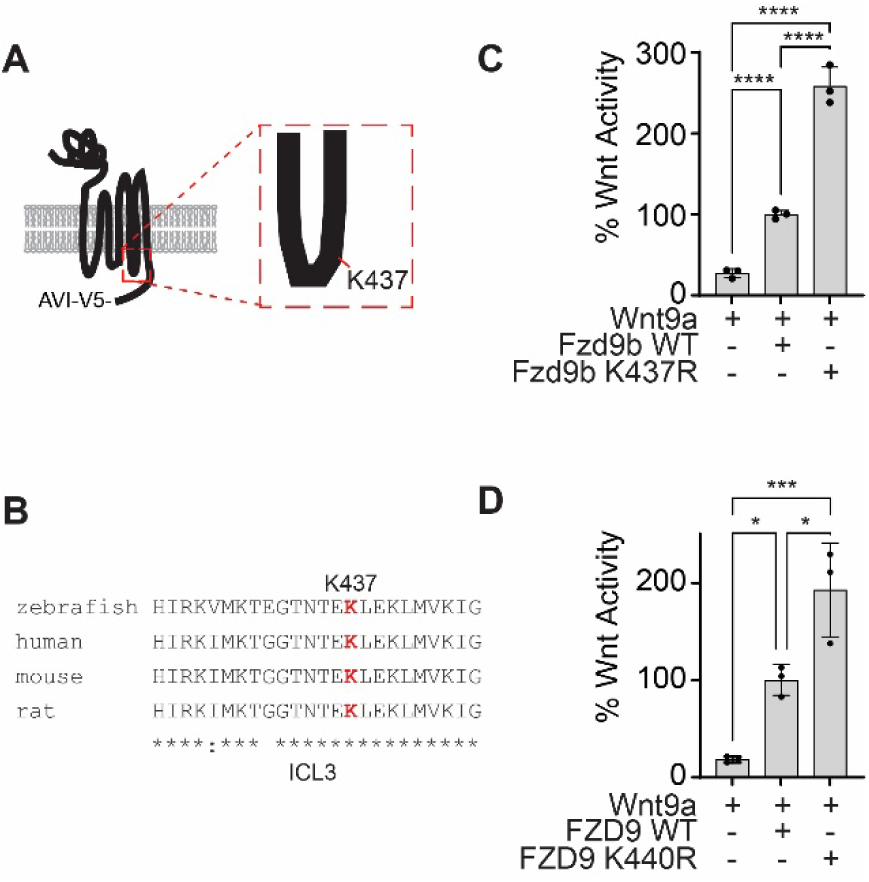
Ubiquitination site at K437 of Fzd9b is required to dampen the Wnt9a signal. **A.** Fzd9b diagram with the K437 ubiquitination site annotated in the third intracellular loop. **B.** Clustal Omega alignment of third intracellular loop (ICL3) peptide sequences from species as noted. Note the high degree of conservation, including at K437. **C.** Super TOP Flash assays to measure Wnt responses in HEK293 cells transfected with zebrafish Fzd9b, Fzd9b^K437R^, and Wnt9a as indicated. **D.** Super TOP Flash assays to measure Wnt responses in HEK293 cells transfected with human FZD9, FZD9^K440R^, and WNT9A as indicated. All data presented are from N=3 biological replicates, with different experiments conducted on different days a total of 3 times with similar results. *P<0.05, ***P<0.001, ****P<0.0001 by ANOVA with Tukey post-hoc comparison.

### Trip12-mediated ubiquitination dampens the Wnt9a/Fzd9b signal

By modeling Wnt signals in human (e.g., HEK293) cells with zebrafish proteins, we can uncover conserved molecular mechanisms of signal regulation^11–13^. Through APEX2-mediated proximity ligation in HEK293 cells, we previously identified human proteins enriched at the intracellular regions of zebrafish Fzd9b in response to zebrafish Wnt9a^11^. From this dataset, we identified ubiquitin ligases that were associated with Fzd9b, and found that the E3 ubiquitin ligase Thyroid hormone Receptor Interacting Protein 12 (TRIP12) had the highest increase in enrichment upon Wnt9a stimulation, followed by other positive regulators of ubiquitination including MYC binding protein 2 (MYCBP2), MSL complex subunit 2 (MSL2), RB binding protein 6 (RBBP6), and UBA domain containing 2 (UBAC2) (Fig. 3A). Using siRNAs targeted to each of these human mRNAs, paired with a Super TOP Flash assay, we found that suppression of only two of these targets, TRIP12 or MYCBP2, led to an increase in Wnt9a/Fzd9b signaling (Fig. 3B, Supplementary Fig. 3A), suggesting that these play a role in dampening the Wnt9a/Fzd9b signal. TRIP12 was prioritized for follow-up analysis as it was the E3 Ubiquitin ligase most dynamically recruited to Fzd9b in response to Wnt9a (Fig. 3A), and it had the strongest impact on Wnt9a/Fzd9b activity (Fig. 3B, Supplementary Fig. 3B).

**Figure 3:**
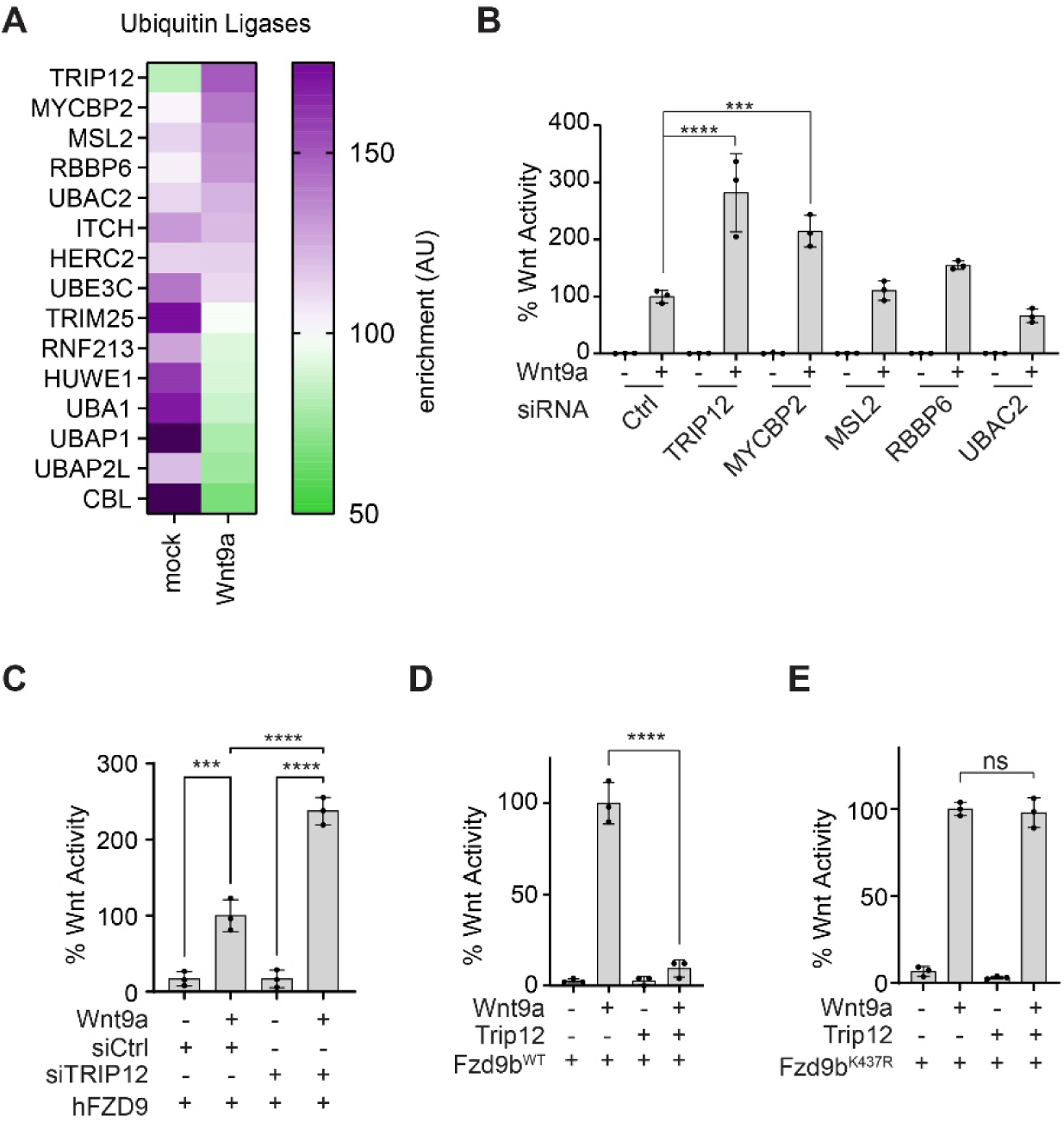
Trip12 dampens the Wnt9a/Fzd9b signal. **A.** Heatmap showing enrichment of ubiquitin ligases recruited to the C-terminal of Fzd9b in response to Wnt9a. Data collected from Grainger, et al, 2019^11^. **B.** Super TOP Flash assays to measure Wnt responses in Fzd9b-mKate HEK293 cells treated with Wnt9a conditioned medium and siRNAs targeted to human mRNAs as indicated. **C.** Super TOP Flash assays to measure Wnt responses in FZD9 HEK293 cells transfected with Wnt9a and siRNAs targeted to human mRNAs as indicated. Super TOP Flash assays to measure Wnt responses in HEK293 cells transfected with Wnt9a, Trip12 and Fzd9b-mKate (**D**) or Fzd9b^K437R^-mKate transfected **(E**). All data presented are from N=3 biological replicates, with different experiments conducted on different days a total of 3 times with similar results. ns-not significant, ***P<0.001, ****P<0.0001 by ANOVA with Tukey post-hoc comparison.

Knockdown of TRIP12 also led to an increase in human FZD9 signaling activity (Fig. 3C), supporting conservation of this function. We found that overexpression of Trip12 nearly ablated the Wnt9a/Fzd9b signal (Fig. 3D), while Trip12 did not impact Fzd9b lacking the putative ubiquitination site (Fzd9b^K437R^) activity, indicating that K437 is the site of Trip12 function (Fig. 3E). Trip12 is an E3 ubiquitin ligase with a highly conserved catalytic cysteine at C2020 (C1959A in humans, Supplementary Fig. 3C), which when substituted for by an alanine ablates ubiquitin ligase activity^20^. Consistent with this, Trip12^C2020A^ did not impact Wnt9a/Fzd9b signaling (Supplementary Fig. 3D), indicating that Trip12 ubiquitin ligase activity impacts Fzd9b function. Altogether, these results demonstrate that Trip12 acts to limit the Wnt9a/Fzd9b activation through ubiquitination of K437 on Fzd9b.

### TRIP12-mediated ubiquitination of Fzd9b at K437 leads to its lysosomal degradation

The E3 ubiquitin ligases RNF43 and ZNRF43 have been shown to broadly regulate the cell surface availability of FZD receptors by targeting them for lysosomal degradation^7–9^, leading us to hypothesize that Trip12 may impact Fzd9b expression. To study Fzd9b cell surface localization, we used our previously established stable Fzd9b-mKate^13^ cells and introduced Trip12 with a C-terminal HA tag. We found that exogenous Trip12-HA led to the loss of Fzd9b expression while the ubiquitin ligase dead Trip12^C2020A^-HA did not (Fig. 4A,B). These data indicated that the E3 ubiquitin ligase activity of Trip12 at K437 decreases Fzd9b availability for signaling.

**Figure 4:**
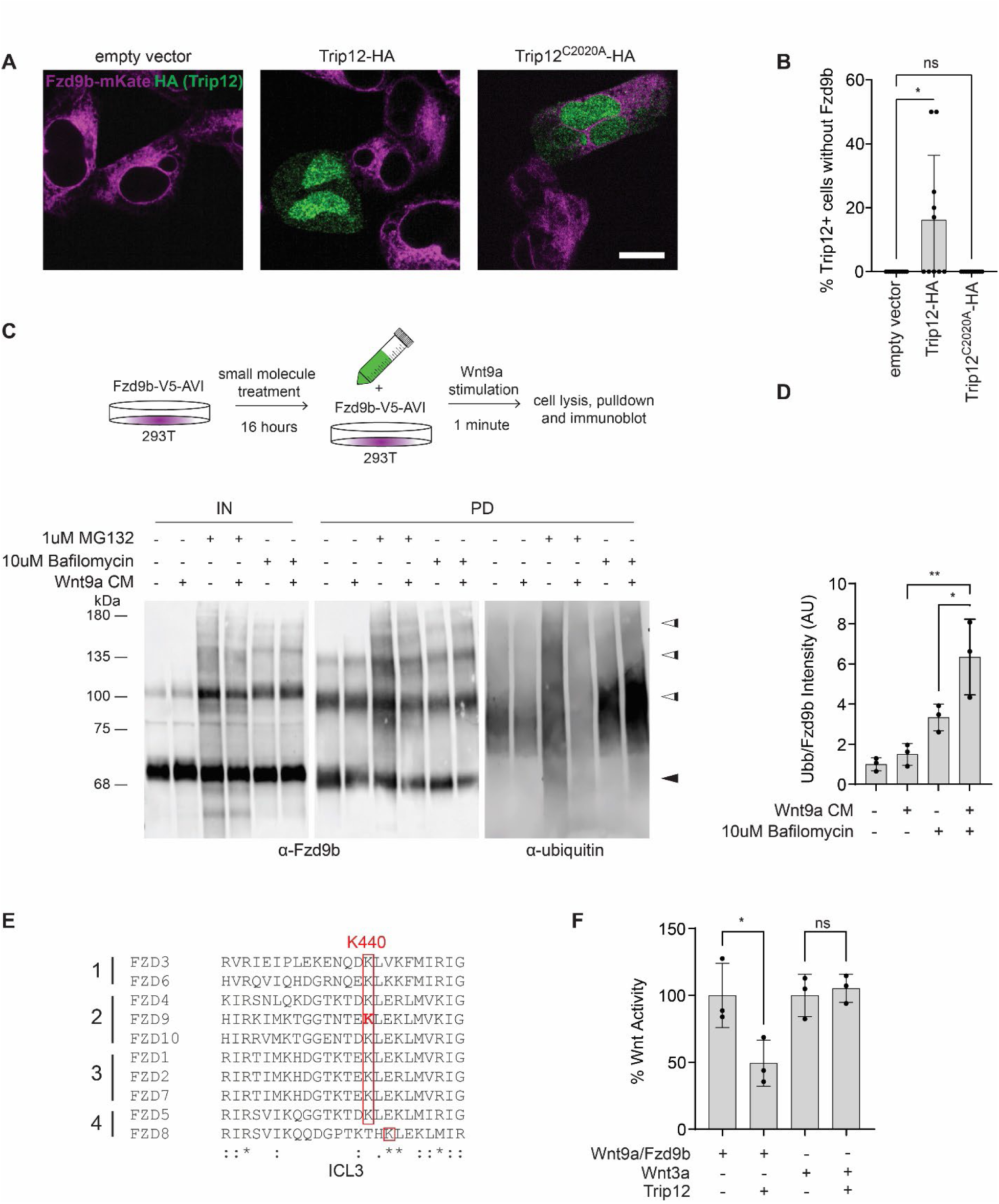
Fzd9b is targeted for lysosomal degradation by Trip12. **A.** Representative confocal Z-plane of stable Fzd9b-mKate cells transfected with empty vector, WT Trip12-HA, or Trip12^C2020A^-HA. Scale bar represents 10 µm. **B.** Ten images from each condition shown in **A** were quantified to determine the percentage of Trip12+ cells that expressed Fzd9b. Three different experiments were conducted on different days with similar results. **C.** Fzd9b-V5-AVI cells were pre-treated with vehicle, 1 µM MG132 or 10 µM BafilomycinA1 for 16 hours, and stimulated with Wnt9a or mock conditioned medium (CM) for 1 minute, lysed, Fzd9b purified with streptavidin beads and immunoblotted. Immunoblots for Fzd9b and ubiquitin antibodies are shown. Black arrowheads indicate the predicted size of unmodified Fzd9b; white arrowheads indicate multiple higher molecular weight forms. **D.** Quantification of PD experiments in **C**, where ubiquitin blot signal was normalized to Fzd9b blot signal in triplicate experiments. **E.** Clustal Omega alignment of third intracellular loop (ICL3) peptides across the 4 human FZD subfamilies. Note the conservation of K440 (equivalent to zebrafish K437). **F.** Super TOP Flash assays to measure Wnt responses in HEK293 cells transfected with Wnt9a/Fzd9b, Wnt3a and Trip 12 as indicated. All data presented are from N=3 biological replicates, with different experiments conducted on different days a total of 3 times with similar results. ns-not significant, *P<0.05, **P<0.01 by ANOVA with Tukey post-hoc comparison.

Ubiquitination of proteins can lead to diverse intracellular outcomes, including degradation of proteins through either the proteasomal or lysosomal pathways^21^. The small molecules MG132 and BafilomycinA1 inhibit the proteasome and lysosomes, respectively^22,23^. Treatment of Fzd9b-V5-AVI cells with MG132 or BafilomycinA1, followed by Wnt9a treatment, V5-tag immunoprecipitation, and subsequent immunoblotting showed an increase in higher molecular weight Fzd9b following treatment with either compound (Fig. 4C), suggesting that both the lysosome and proteasome normally degrade Fzd9b. In addition, there was an increase in the ubiquitinated proteins coinciding with the 110 kDa Fzd9b bands, especially in response to BafilomycinA1, compared to vehicle treatment (Fig. 4C), suggesting that ubiquitinated Fzd9b is degraded primarily by the lysosome. We observed a further increase in ubiquitinated Fzd9b with Wnt9a treatment (Fig. 4C,D), indicating that adding Wnt9a to activate the system pushes Fzd9b toward the lysosome. We verified that the addition of MG132 led to an increase in β-catenin, as previously reported^24^ (Supplementary Fig. 4A), indicating the modest impact noted on Fzd9b accumulation was not due to poor MG132 activity.

Loss of function mutations in other E3 ubiquitin ligases targeting Fzds, such as ZNRF43 and RNF3, are tumorigenic^7,9,10^. In a broad survey of human cancer samples across The Cancer Genome Atlas, we identified 69 different DNA mutations leading to premature STOP codons in the TRIP12 coding sequence, all of which are predicted to ablate the E3 ubiquitin ligase activity (Supplementary Fig. 4B). These mutations were present in every cancer type examined, suggesting that TRIP12 modification of FZD activity may play a broader role in cancer than previously appreciated. Consistent with this, aligning the residues of the third intracellular loop of all zebrafish Fzd proteins revealed that of the 4K residues, only K437 in zebrafish (or K440 in humans) is conserved across all human FZDs (Fig. 4E) and zebrafish Fzds (Supplementary Fig. 4C), suggesting that TRIP12 could impact Wnt signaling broadly. Contrary to this, however, Wnt3a, which is known to signal through most FZD receptors^25^, is unaffected in the presence of overexpressed Trip12 as assessed by Super TOP Flash activity (Fig. 4F). These data indicate that there is some degree of specificity for TRIP12-mediated ubiquitination of Fzd9b in dampening the Wnt9a signal.

### Trip12 is required for hematopoietic stem cell development

The pairing of Wnt9a and Fzd9b is required for hematopoietic stem cell proliferation in the aorta in zebrafish, which can be seen by a loss of *cmyb*+ cells at 40 hours post fertilization (hpf) in the absence of the Wnt9a signal^11,12^. We hypothesized that a loss of Trip12 would lead to an increase in Fzd9b receptor availability and an increase in Wnt9a/Fzd9b signaling, ultimately leading to an increase in *cmyb+* cells at 40 hpf. Injection of a translation blocking (ATG) antisense morpholino (MO) targeting *trip12* showed an increase in *cmyb* expression at 40 hpf (Fig. 5A), suggesting an increase in hematopoietic stem cells. After hematopoietic stem cells emerge from the dorsal aorta, they proceed to seed the caudal hematopoietic tissue, where we observed a rise in *cd41*^+^ (Fig. 5B, C)^26^ and *gata2b^+^* (Supplementary Fig. 5)^27^ hematopoietic stem and progenitor cells in MO treated animals at 72 hpf (Fig. 5B)^26^. The MO was rescued by *trip12* mRNA, indicating specificity of the treatment (Supplementary Fig. 5). Altogether, these data support a role for Trip12 in hematopoietic stem cell development.

**Figure 5:**
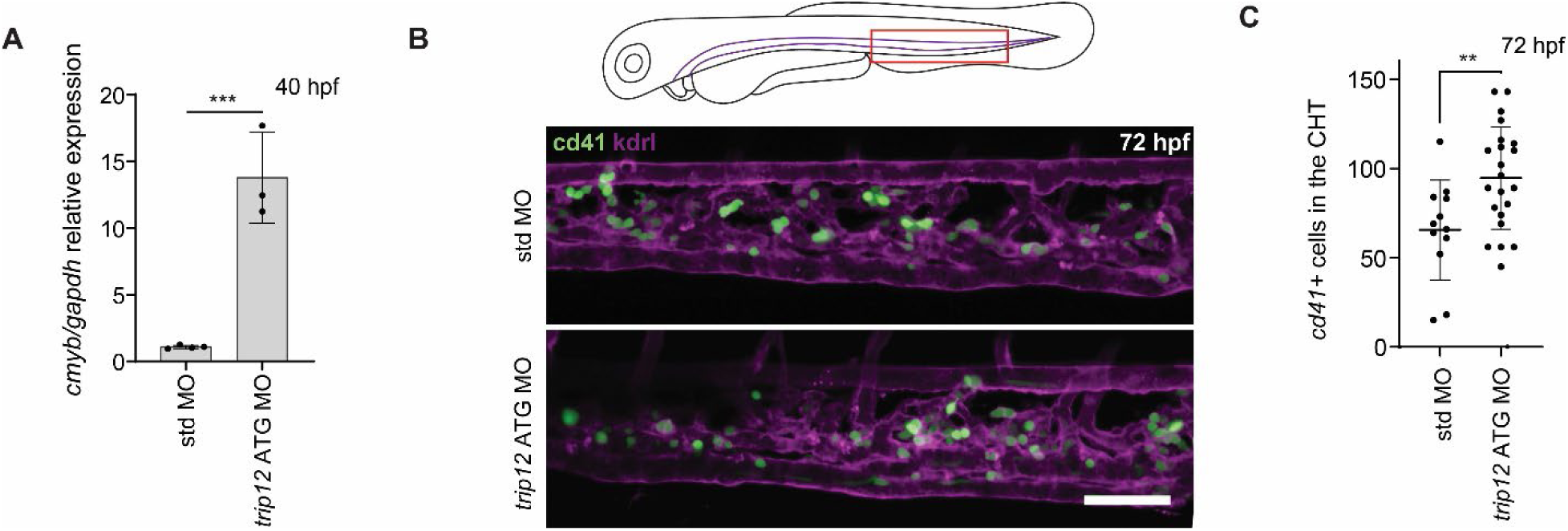
Trip12 regulates hematopoietic stem cell development. **A.** Zebrafish at 40 hpf, injected with either standard (std) or *trip12* ATG morpholinos (MO), hematopoietic stem and progenitor cell maker *cmyb* quantified by qPCR. N=3 biological replicates are shown. ***P<0.001 by ANOVA with Tukey post-hoc comparison. **B.** Representative confocal Z-stacks of the caudal hematopoietic tissue of 72 hpf *cd41:eGFP* (green); *kdrl:mCherry-CAAX* (magenta) zebrafish injected with either std or *trip12* ATG MO. Scale bar represents 50 µm. **C.** Quantification of hematopoietic stem and progenitor cells marked by GFP in **B**. Each dot represents a biological replicate. **P<0.01 by ANOVA with Tukey post-hoc comparison.

## Discussion

Tight regulation of Wnt signaling is critical during development and homeostasis, partly due to its important role in cellular proliferation. For example, we have shown that Wnt9a and Fzd9b are essential for developmental proliferation of hematopoietic stem cells^11,12^. The Fzd9b receptor is available at the cell membrane without a Wnt ligand but is highly activated in the presence of Wnt9a. Here, we demonstrate that Trip12 is required to limit the Wnt9a/Fzd9b signal during hematopoietic stem cell development through its E3 ubiquitin ligase activity. Trip12 ubiquitinates Fzd9b in the third intracellular loop at K437, leading to its lysosomal targeting and degradation, thus limiting Fzd9b cell surface expression available for pathway activation. Wnt9a stimulation brings together the Fzd9b complex, allowing β-catenin accumulation in the cytoplasm, and translocation to the nucleus, where it activates target genes required for hematopoietic stem cell proliferation (Fig. 6).

**Figure 6:**
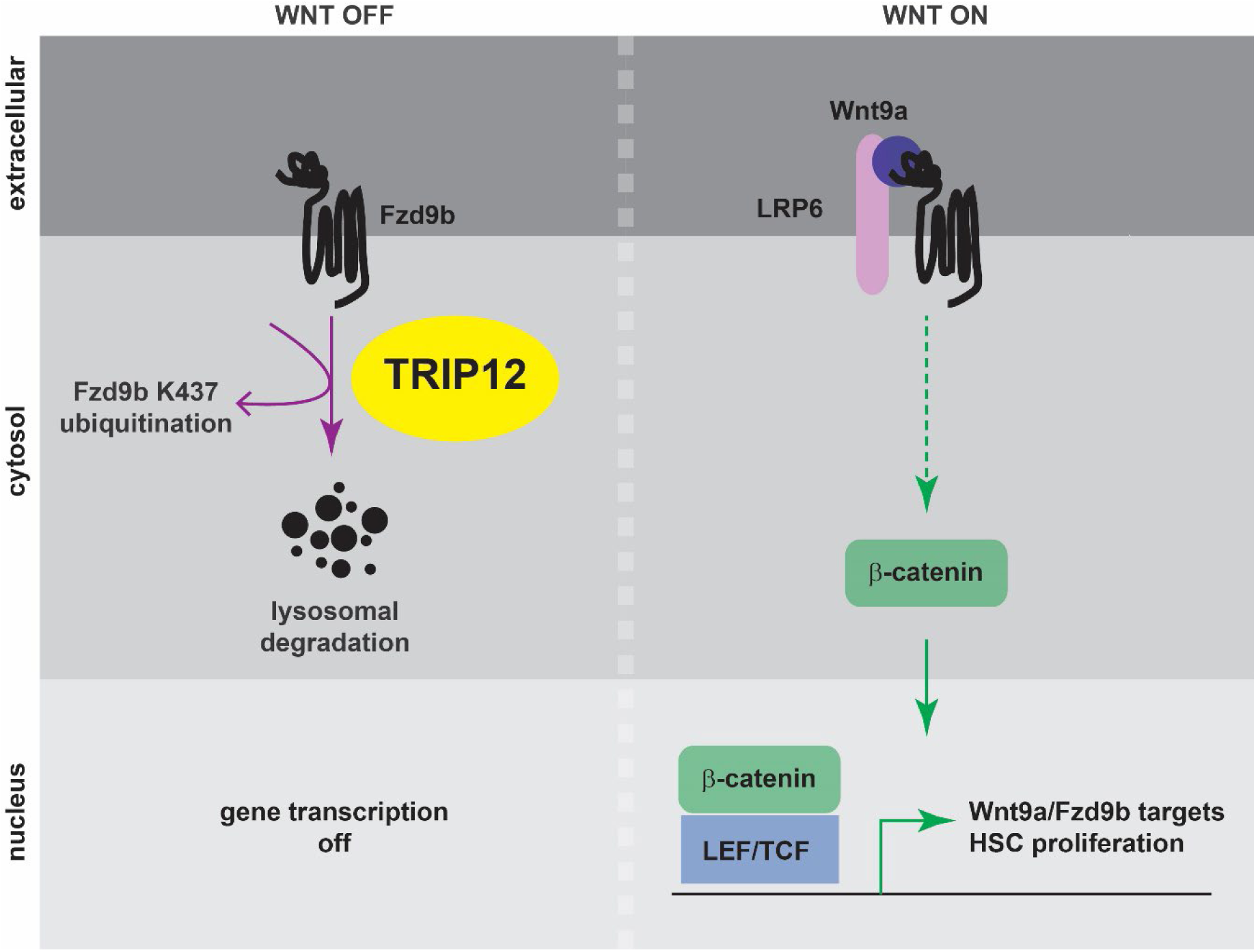
A model for Trip12 regulation of Fzd9b and hematopoietic stem cell development. In the absence of Wnt9a, Fzd9b is ubiquitinated at K437, leading to its lysosomal degradation. Wnt target gene activation is shut off. When Wnt9a is present, the Fzd9b receptor complex, including LRP5/6 is assembled, leading to β-catenin translocation to the nucleus, and activation of Wnt9a/Fzd9b target genes that drive hematopoietic stem cell proliferation.

There has been considerable debate in the Wnt field about the positive and negative influences of endocytosis on signaling^28–40^. These differences may reflect specific cell and receptor context-dependent requirements for internalization of the Wnt/Fzd complex as a pre-requisite for signaling^13,37^, as opposed to formation of signalosomes^41^, or trafficking of co-receptors without a ligand^42,43^. In addition, many of these observed differences may lie in the regulation of receptor availability at the membrane, as we describe here.

The balance of ubiquitination and deubiquitination has been proposed to regulate Wnt signaling sensitivity to cells through trafficking Fzd receptors to the lysosome or the cell membrane, as demonstrated with Fzd4^44^. Other groups have shown the importance of different ubiquitin ligases in regulating Wnt signals through downregulating cell surface expression of different FZD receptors. For example, RNF43 regulates FZD1, FZD5 and FZD7 cell surface expression^9^, while ZNRF3 regulates FZD4, FZD5, FZD6 and FZD8^7^, with a preference for FZD6^45^. Mechanistically, RNF43 and ZNRF3 function to ubiquitinate cytosolic lysine residues^9^, a process that is inhibited by introducing the secreted ligand R-Spondin, which acts in concert with the seven-pass transmembrane LGR4/5 receptors^7^. The regulation of FZD cell surface expression by RNF43 and ZNRF3 is thought to occur through the lysosome^9,10,44^. Our findings here show that TRIP12 regulates Fzd9b/FZD9 cell surface expression, through ubiquitination of a single intracellular lysine residue in the third intracellular loop. However, this mechanism is distinct from the transmembrane receptors RNF43 and ZNRF3, since TRIP12 is located intracellularly. TRIP12 is enriched to Fzd9b upon Wnt9a stimulation (Fig. 3A), suggesting that there is a Wnt9a-dependent enhancement of Fzd9b regulation. This is further supported by the enhancement of ubiquitinated Fzd9b in response to Wnt9a stimulation, especially in the presence of BafilomycinA1 (Fig. 4C,D). Interestingly, we observed that the K437 ubiquitination site is conserved across all FZDs (Fig. 4E, Supplementary Fig. 4C), yet Wnt3a (which signals through most FZDs) signaling activity, was not impacted by Trip12 (Fig. 4F). This may suggest additional layers of regulation required for Fzd9b-specific ubiquitination by Trip12. For example, we previously showed that EGFR phosphorylation of the Fzd9b C-terminal tail is required to initiate endocytosis and Wnt9a signaling. It is tempting to speculate that such modifications are a pre-requisite for Trip12 function on Fzd9b as well.

We observed that Wnt9a does not impact overall Fzd9b abundance (Fig. 1C, Fig. 4C). However, the ubiquitinated pool of Fzd9b is increased with Wnt9a treatment (Fig. 4C,D), and Fzd9b is lost in the presence of Trip 12 (Fig. 4A,B). These data support a model where only a portion of the Fzd9b pool is ubiquitinated at a given time. Loss of the ubiquitination site at K437 leads to a 3-fold increase in Wnt9a/Fzd9b signaling activity (Fig. 2B), indicating that although the pool of ubiquitinated Fzd9b is limited, there is an impact on function. In addition, when comparing the pool of ubiquitinated Fzd9b regulated by the proteasome (MG132, Fig. 4C) to the pool regulated by the lysosome (Bafilomycin, Fig. 4C), it appears as though more ubiquitinated Fzd9b is processed through the lysosome than the proteasome. This may indicate that there are different pools of Fzd9b; some may be signaling competent, while others may not. This also may reflect differences in signaling requirements across tissues, and Fzd receptors. For example, in *Drosophila* imaginal discs, Frizzled2 is sent for degradation only after a signal-competent complex with Arrow (equivalent to LRP6) is formed^46^. Additionally, there are tissue specific differences in degradation routes and timing of Drosophila Frizzled2^31^. Although we did not detect other ubiquitination sites on Fzd9b in this study, this may be related to relative instability and/or abundance of peptides targeted with ubiquitination.

TRIP12 function is incompletely understood, although its dysregulation is implicated in multiple diseases^47^. TRIP12 promotes proteolysis of at least nine substrates, including chromatin regulators^48,49^, ubiquitin machinery^50,51^, and transcription factors^20,52^, among others^53–55^. In addition to these *bona fide* substrates, TRIP12 has been noted to interact with a huge swath of proteins in the cell, including several receptors; however, targeting these interacting proteins has yet to be shown^47^. Interestingly, TRIP12 is reported to be localized in the nucleus in cell lines and human tissues^51,56^. Although we found a predominant nuclear signal, Trip12 was also expressed outside the nucleus (Fig. 4A). Consistent with its hypothesized role in regulating diverse cellular substrates, mutations in *TRIP12* are commonly identified in cancers. Notably, premature STOP codons identified (Supplementary Fig. 4B) would all be predicted to impact ubiquitin ligase function. Our findings here show that TRIP12 impacts Wnt9a/Fzd9b signaling now link TRIP12 function to the regulation of the WNT pathway, which is also widely implicated in cancer initiation and progression^1,57–60^. It has been challenging to generate therapeutics targeting the WNT pathway due to its pleiotropic requirements in homeostasis. Identifying specific WNT pathways in vivo will be essential to derive therapies that more specifically target cancer cells. The interaction between TRIP12 and FZD9 may represent one such avenue for therapeutic intervention.

## Materials and Methods

### Cell culture

All luciferase assays were conducted in cell lines derived from our previously established HEK293 cells with a stably integrated Super-TOP-Flash reporter (STF)^18^. All cells were grown under standard conditions with media supplemented with 10% heat-inactivated fetal bovine serum (FBS) and 1% penicillin/streptomycin. HEK293 cells used Dulbecco’s modified Eagle’s medium (DMEM) as a base, while Chinese Hamster Ovary (CHO) cells were grown in a 50:50 mix of DMEM and F12. Mycoplasma testing was conducted monthly, and cells were always negative.

### Generation of transgenic cell lines and Wnt conditioned medium and luciferase reporter assays

Antibiotic-selected clonal transgenic cell lines were generated as previously described^11^, using the PiggyBAC transposase system^61^. Clonal lines were validated using immunoblotting and Wnt reporter assays.

Wnt protein-containing conditioned media (including Wnt9a-Gam) was prepared from transgenic CHO cell lines as previously described ^13^.

Luciferase reporter assays were performed with either transient transfection, or stable lines, as indicated in the figure legends, and as previously described^13^. Each experiment was performed with at minimum biological triplicate samples and reproduced at least once with a similar trend. Wnt activity was calculated by normalizing Firefly Luciferase output to Renilla Luciferase; Wnt9a/Fzd9b fold induction was set to 100. siRNA sequences used are available as a Supplementary Table 1.

### Fzd9b protein enrichment and immunoblotting

Enrichment of Fzd9b proteins was accomplished using HEK293 cells stably expressing V5 and Avi tagged zebrafish Fzd9b, and co-expressing BirA after a P2A ribosomal skip sequence. Cells were grown in standard media as described above, supplemented with 0.2mg/L biotin (Sigma, B4639) for at least 24 hours. Cells were treated as described in the figure legends and lysed in RIPA buffer (150mM NaCl, 50mM Tris-HCl, pH7.5, 1% NP40, 0.1% SDS, 12mM deoxycholate) supplemented with protease inhibitor cocktail (Roche, 04693132001), phosphatase inhibitor cocktail (Thermo, A32957) and 20mM N-Ethylamaleimide (Fisher, E01365G). Biotinylated proteins were enriched using Dynabeads MyOne Streptavidin T1 (Fisher, 65601), essentially according to the manufacturer’s recommendations, including five 1mL washes with ice cold RIPA buffer. Biotinylated proteins were eluted from the beads by resuspending in Laemmli buffer^62^, supplemented with 25mM biotin-D and incubating at 55°C for 15 minutes.

Immunoblots were performed according to LiCor manufacturer’s procedures (lysates subjected to blotting for Fzds were not boiled), using primary antibodies for: α-V5, [1:5,000, Cell Signaling Technologies, 13202S], a-b-actin, [1:10,000, Sigma, A2228], 1:500, α-Fzd9b^11^, 1:500, α-ubiquitin [BioLegend, 646302, Lot# B287176]. Signals were detected using IRDye 800CW Streptavidin (for biotinylated proteins) [1:25,000, LiCor, 926-322230, Lot#D20920-05], goat-a-mouse IR700 [1:20,000, LiCor, 926-68070, Lot#D20419-25] and goat-α-rabbit IR800 [1:20,000, LiCor, 926-32211, Lot#D20510-25].

### Protein digestion and sample preparation

Fzd9b was eluted from streptavidin beads using 25mM biotin D, 2.5% SDS and 5% beta-mercaptoethanol at 95℃ for 5 minutes, and proteins were resolved by SDS PAGE and stained using EZBlue (Fisher, G1041), according to the manufacturer’s recommendations. The gel band was tryptic digested with Tryptic Digestion Kits according to the manufacturer’s recommendations (Thermo Fisher, 89871). The supernatants were dried down on SpeedVac and resuspended with 10 µl of 0.1% trifluoroacetic acid in H_2_O. Synthetic peptides for TEGTNTEK(GG)LEK were obtained from Biosynth (Gardner, MA).

### LC-MS/MS for Global and Targeted Analysis

Samples were analyzed on an Exploris 480 (Thermo Scientific) attached to a Vanquish Neo nano UPLC (Thermo Scientific). Peptides were separated with nano HPLC column (75 μm x 20 cm, 1.7 µm C18, P/N HEB07502001718IWF, CoAnn Technologies). Mobile phase A was water with 0.1% formic acid. The LC gradient was as follows: 1% B (20% water, 80% acetonitrile, 0.1% formic acid) to 26% B over 51 minutes, 85% B over 5 minutes, and 98% B over 4 minutes, with a total gradient length of 60 minutes. One full MS scan was collected with a scan range of 350 to 1200 m/z, resolution of 120 K, standard maximum injection time, and Auto Gain Control (AGC). A series of MS2 scans were acquired for the most abundant ions from the MS1 scan using orbitrap analyzer. Ions were filtered with charge 2–5. An isolation window of 1.6m/z was used with quadruple isolation mode. Ions were fragmented using higher-energy collisional dissociation (HCD) with a collision energy of 30%. For the targeted quantitation, parallel reaction monitoring (PRM) was performed on an Exploris 480 mass spectrometer coupled with the Vanquish Neo LC system (Thermo Fisher Scientific). A total of 2 μg of digested peptides were separated on a nano capillary column (20 cm × 75 μm I.D., 365 μm O.D., 1.7 μm C18, CoAnn Technologies, Washington) at a flow rate of 300 nL/min. Gradient conditions were identical to above. Full MS spectra (m/z 375-1200) were collected at a resolution of 120,000 (FWHM), and MS2 spectra at 30,000 resolutions (FWHM). Standard AGC targets and automatic maximum injection times were used for both full and MS2 scans. A 32% HCD collision energy was used for MS2 fragmentation. All samples were analyzed using a multiplexed PRM method with an unscheduled inclusion list containing the target peptide TEGTNTEK(GG)LEK.

### Data Analysis

Proteome Discoverer 3.1 (Thermo Scientific) is used to process raw spectra. The search criteria are protein database; Danio rerio (Taxonomy ID: 7955). Carboxyamidomethylated (+57 Da) at cysteine residues for fixed modifications, GlyGly at lysine (+114 Da), oxidized at methionine (+16 Da) residues, and acetylation at lysine residues (+14 Da) for variable modifications, two maximum allowed missed cleavage, 10 ppm tolerance for MS and MS/MS. All PRM data analysis and integration were performed using Skyline software^63^. The transitions’ intensity rank order and chromatographic elution were required to match those of synthetic standards for each measured peptide.

### Plasmids

Expression constructs were generated by standard means using PCR from plasmids harboring a CMV promoter, or a doxycycline inducible promoter system. Fzd9b and FZD9 plasmids were previously described^11,13^. Mutations in Fzds were introduced using site directed mutagenesis. Trip12 cDNA (NM_001347668.1) was PCR amplified from zebrafish cDNA. AVI Tag (GLNDIFEAQKIEWHE) coding sequence was inserted via site-directed mutagenesis. BirA (R118G) was synthesized by Integrated DNA Technologies. All plasmids were sequence validated by full-plasmid sequencing.

### Confocal imaging and analysis for Wnt9a/Fzd9b trafficking

Confocal imaging and analysis for Wnt9a/Fzd9b trafficking were done essentially as previously described ^13^. Briefly, cells were grown and fixed on glass coverslips stained by immunofluorescence using the primary antibodies (α-V5, [Cell Signaling Technologies, 13202S], α-Fzd9b^11^, α-LAMP1, [Cell Signaling Technology, cat# 15665S]), and secondary antibodies (1:500 for each, Goat-α-mouse Alexa 488 [cat# A11001, lot# 2189178], goat-α-rabbit Alexa 488 [cat# 4050-30, lot# e1317 NC39], goat-α-mouse Alexa 555 [cat# 1030-32, lot# E2518 ZA08E], goat-α-rabbit Alexa 647 [cat# 4050-31, lot# E1519 S109Z], Southern Biotech). Slides were imaged within one week.

Confocal z-slices were collected using a Andor Dragonfly 620-SR spinning disc mounted on a Leica DMi8 microscope, equipped with 405, 488, 561, 640 and 730 nm laser lines and 5x air, 10x air, 20x air, 40x water, 63x oil and 100x oil objectives. Scanning was sequential and images were collected at a minimum of 1900 x 1900 resolution. Representative images are single z-slices. Calculation of % events was performed by examining z-stacks through the entire cell and identifying stable Fzd9b-mKate cells that were positive for HA-tagged Trip12 constructs.

### Zebrafish husbandry and mRNA rescue

Zebrafish were maintained and propagated according to Van Andel Institute and local Institutional Animal Care and Use Committee policies. AB* zebrafish were used as wild-type animals in all experiments. The Tg(cd41:eGFP)^la2Tg^, Tg(gata2b:KalTA4)^sd32Tg^, Tg(UAS:GFP)^mu271^ and Tg(kdrl:mCherry-CAAX)^y171^ lines have been previously described^27,64,65^. The ATG MO for *trip12* with the sequence 5’-GACATTGGCACCTCTCTCCTGAAG-3’ was acquired from GeneTools. 1-cell stage zygotes were injected with ATG-blocking *trip12* MO at dosages indicated in the figures. Embryos and larvae were cultured to the ages indicated in figures in Essential 3 (E3) medium (5 mM NaCl, 0.17 mM KCl, 0.33 mM CaCl2, 0.33 mM MgSO4, 10^-5^ % Methylene Blue).

The mRNA for rescue experiments was synthesized using the SP6 mMessage machine kit (LifeTech), according to the manufacturer’s recommendations. The resultant mRNA was quantified by nanodrop, and dosages are indicated in figures.

### qPCR analysis

Standard methods were used to derive RNA and cDNA, and qPCR was performed using FastStart Universal SYBR Green Master Mix (Roche) according to the manufacturer’s recommendations and analyzed using the 2^-DDCt^ method^66^. Primers used are available as a Supplementary Table 2.

## Acknowledgements

Mass spectrometry was performed by the Van Andel Institute Mass Spectrometry Core (RRID: SCR_024903); Hyoungjoo Lee and Ryan Sheldon are thanked for data acquisition. Imaging was performed in part in the Van Andel Institute Optical Imaging Core (RRID:SCR_021968). Research reported in this publication was supported by the National Institute of General Medical Science under Award Number R35GM142779 (SG) and R35GM137976 (ANS), and by the National Institute of Cancer under Award Number T32CA251066 (ADI). The content is solely the responsibility of the authors and does not necessarily represent the official views of the National Institutes of Health.

## Author Contributions

JE, ADI, CG, VT, IC and NJL designed, conducted and analyzed experiments, and edited the manuscript. ANS designed experiments, performed analysis, and edited the manuscript. SG conceived, designed, supervised experiments and analysis, and wrote the manuscript.

## Competing Interests

The authors do not report any competing interests.

## Data and materials availability

All materials are available upon reasonable request.

## Correspondence

Correspondence and requests for materials should be addressed to SG at.

---

**Supplementary Figure 1:**
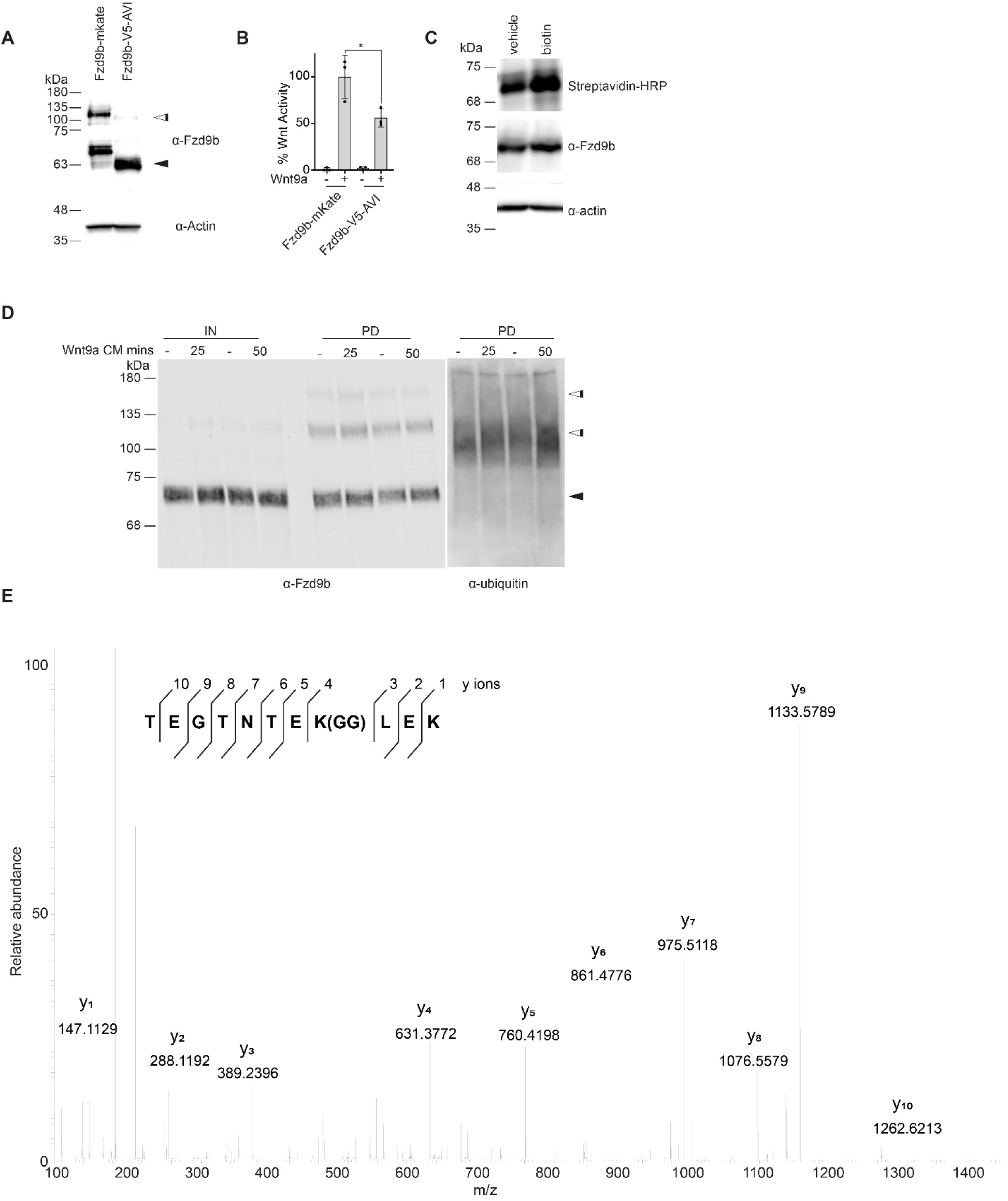
Fzd9b is ubiquitinated. **A.** Representative immunoblot of protein extracted from HEK293 Fzd9b-mKate or Fzd9b-V5-AVI cells. Black arrowhead points to the expected size for Fzd9b; white arrowhead points to additional sized bands detected by the Fzd9b antibody. **B.** Super TOP Flash assays to measure Wnt responses in Fzd9b-mKate or Fzd9b-V5-AVI HEK293 cells treated with Wnt9a conditioned medium as indicated. All data presented are from N=3 biological replicates, with different experiments conducted on different days a total of 3 times with similar results. *P<0.05 by ANOVA with Tukey post-hoc comparison. **C**. Representative immunoblot of protein extracted from HEK293 Fzd9b-V5-AVI cells treated with vehicle or biotin. **D**. Immunoblot of input (5%, IN) and pulldown (PD) of protein extracted from HEK293 Fzd9b-V5-AVI cells treated with mock or Wnt9a conditioned medium (CM) for 25 or 50 minutes as indicated. Blots for Fzd9b and ubiquitin antibodies are shown. Black arrowhead points to the expected size for Fzd9b; white arrowheads point to additional sized bands detected by the Fzd9b antibody. **E**. MS2 spectra of the synthetic ubiquitinated peptide (TEGTNTEK(GG)LEK) of Fzb9b, displaying the relative abundance of the fragment ions. All fragment ions were manually assigned. GG denotes the di-glycyl remnant produced on ubiquitinated lysine residues (K-ε-GG) following trypsin digestion.

**Supplementary Figure 2:**
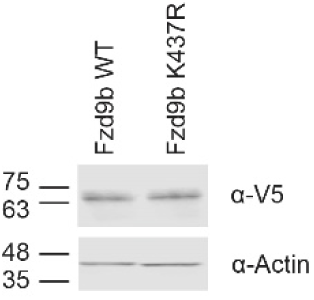
Fzd9b K437R does not impact on Fzd9b expression. Immunoblot of WT and K437R Fzd9b protein lysates.

**Supplementary Figure 3:**
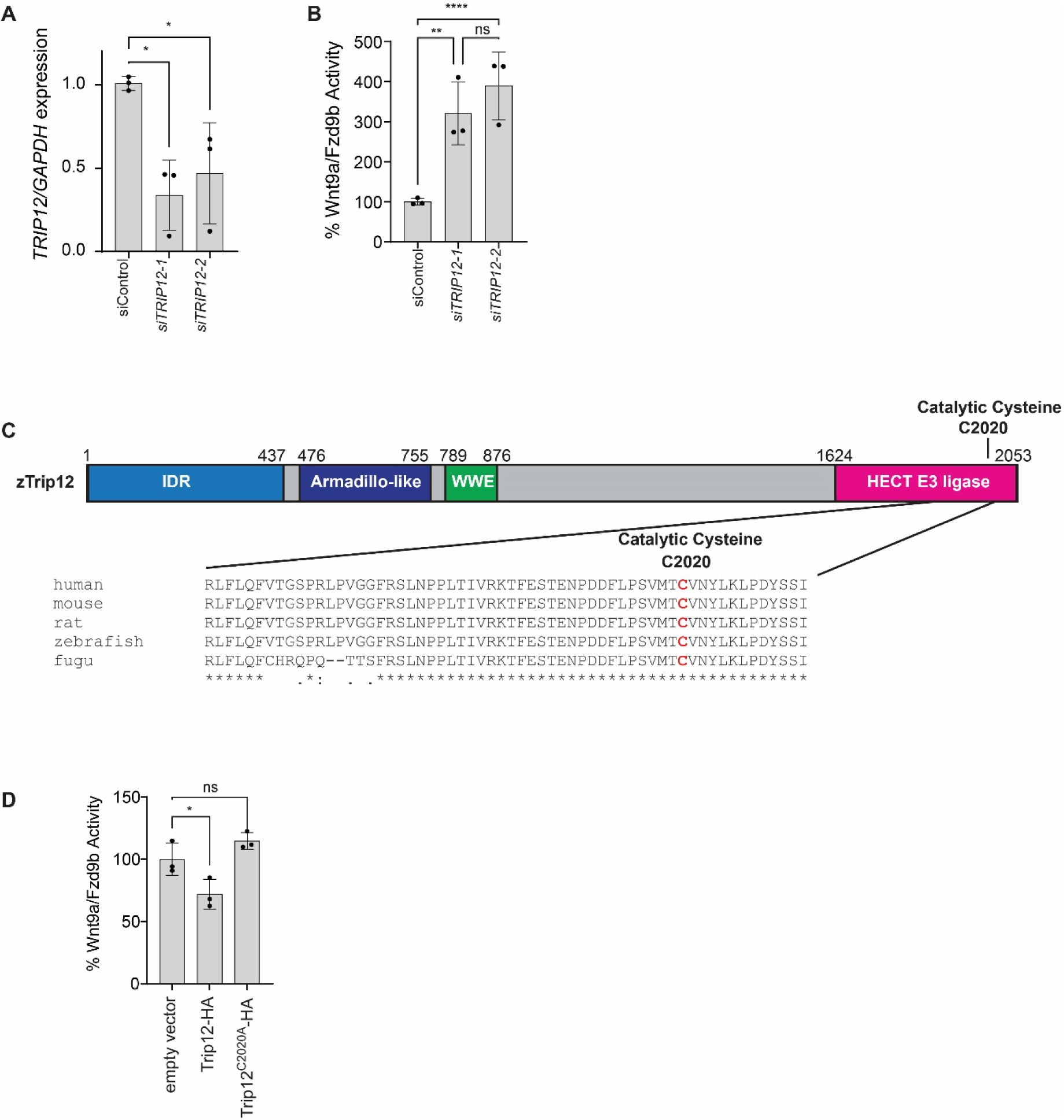
TRIP12 regulates Wnt9a/Fzd9b signaling. **A.** Relative quantification of human *TRIP12* transcript following transfection with control, or 2 different *TRIP12* siRNAs. **B.** Super TOP Flash assays to measure Wnt responses in Fzd9b-V5-AVI HEK293 cells treated with siRNAs and Wnt9a conditioned medium as indicated. **C.** Diagram of zebrafish Trip12 with protein domains noted. Below, alignment of several species of Trip12 showing conserved catalytic cysteine. **D.** Super TOP Flash assays to measure Wnt responses in Fzd9b-V5-AVI HEK293 cells transfected with empty vector, Trip12, or Trip12^C2020A^, and treated Wnt9a conditioned medium as indicated. All data presented are from N=3 biological replicates, with different experiments conducted on different days a total of 3 times with similar results. ns-not significant, *P<0.05, **P<0.01, ****P<0.0001 by ANOVA with Tukey post-hoc comparison.

**Supplementary Figure 4:**
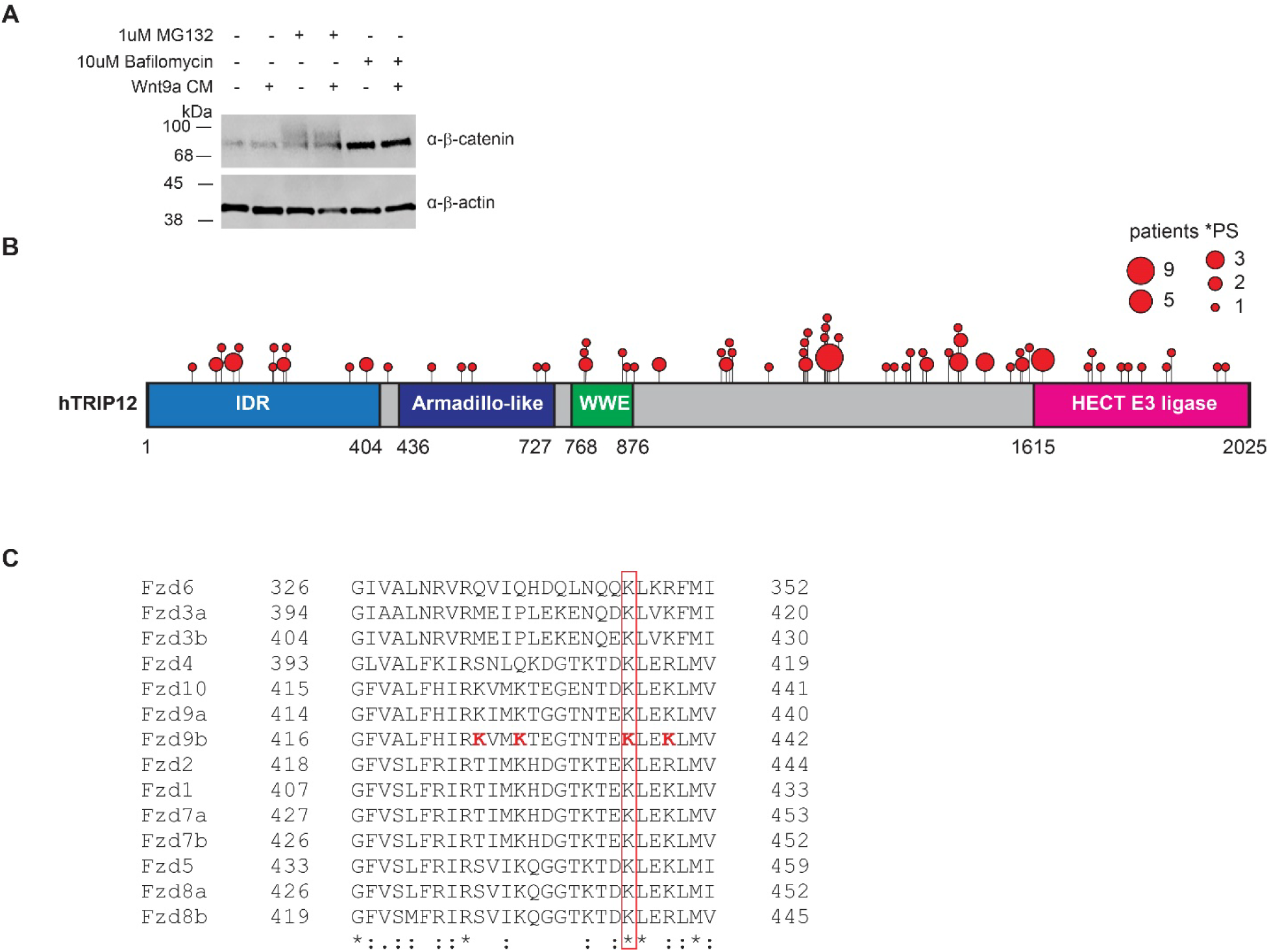
Fzd9b is degraded by the lysosome in response to Trip12. **A.** Immunoblot of protein extracts treated with vehicle, 1 µM MG132 or 10 µM Bafilomycin and Wnt9a conditioned medium (CM) as indicated. **B**. Diagram of mutation frequencies in cancers from data in The Cancer Genome Atlas. Each red spot indicates the location of mutations; the size of red spots indicates the number of patients identified with a mutation. Only nonsense mutations predicted to case premature STOP (PS) codons are shown. **C.** Clustal Omega alignment of third intracellular loop (ICL3) of all zebrafish Fzd proteins showing K437 is conserved.

**Supplementary Figure 5:**
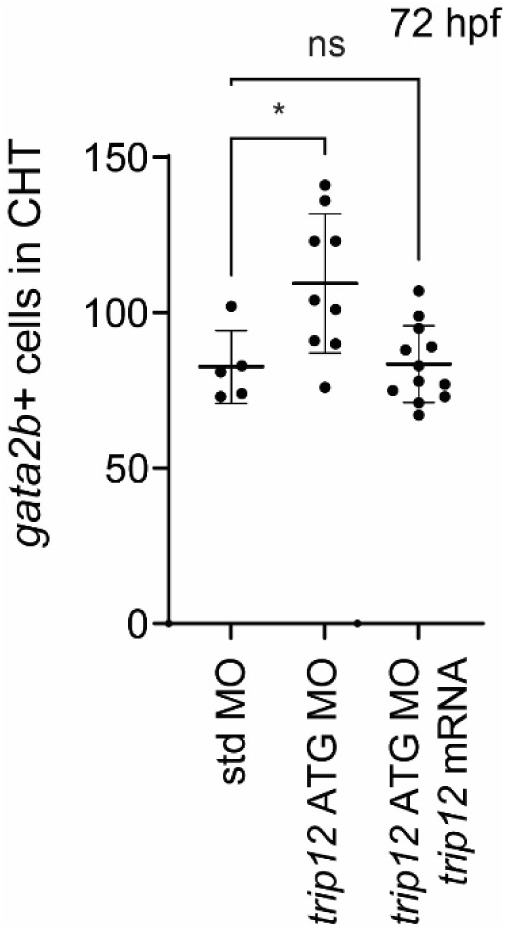
Trip12 is required for hematopoietic stem cells development *in vivo*. Quantification of hematopoietic stem and progenitor cells marked by GFP in Tg(*gata2b:KalTA4;UAS:GFP)* zebrafish caudal hematopoietic tissue at 72 hpf. Each dot represents a biological replicate. *P<0.05 by ANOVA with Tukey post-hoc comparison.

**Supplementary Table 1:**
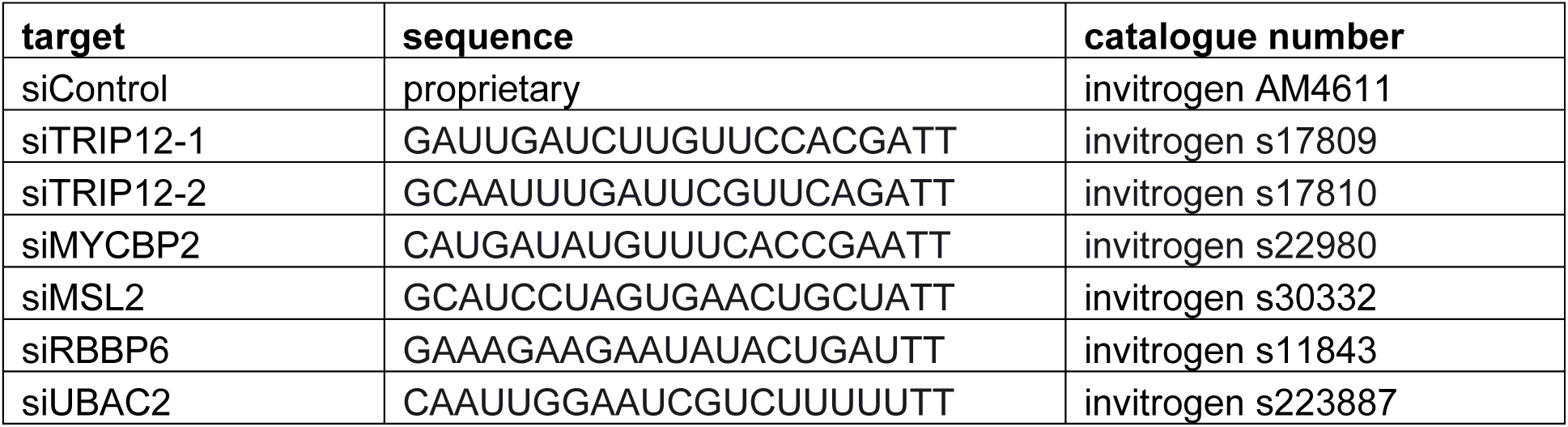
siRNAs used.

**Supplementary Table 2:**
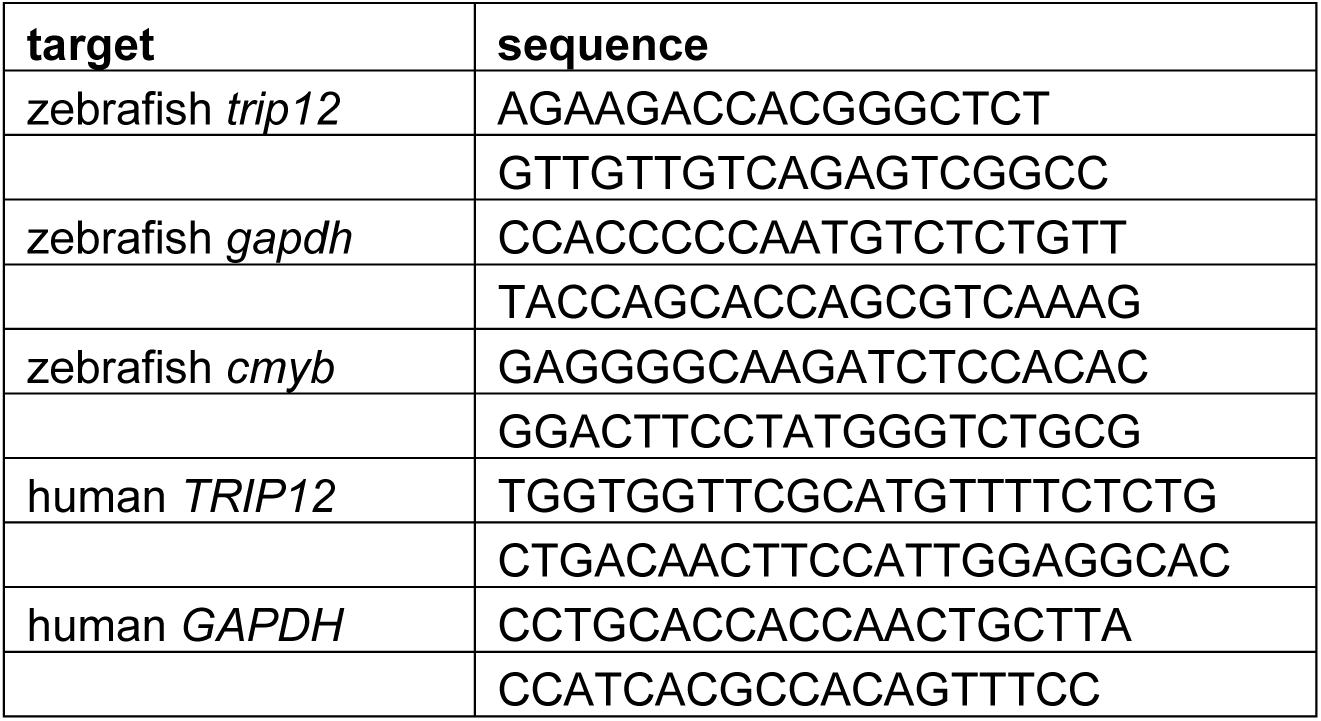
qPCR primers used.

## Notes

### Competing Interest Statement

The authors have declared no competing interest.

